# A novel homology-based algorithm for the identification of physically linked clusters of paralogous genes

**DOI:** 10.1101/051953

**Authors:** Juan F. Ortiz, Antonis Rokas

**Author notes:** To whom correspondence should be addressed: Antonis Rokas, Department of Biological Sciences, Vanderbilt University, VU STATION B #35-1634 Nashville, TN 37235, USA.

## Abstract

Highly diverse phenotypic traits are often encoded by clusters of gene paralogs that are physically linked on chromosomes. Examples include olfactory receptor gene clusters involved in the recognition of diverse odors, defensin and phospholipase gene clusters involved in snake venoms, and Hox gene clusters involved in morphological diversity. Historically, gene clusters have been identified subjectively as genomic neighborhoods containing several paralogs, however, their genomic arrangements are often highly variable with respect to gene number, intergenic distance, and synteny. For example, the prolactin gene cluster shows variation in paralogous gene number, order and intergenic distance across mammals, whereas animal Hox gene clusters are often broken into sub-clusters of different sizes. A lack of formal definition for clusters of gene paralogs does not only hamper the study of their evolutionary dynamics, but also the discovery of novel ones in the exponentially growing body of genomic data. To address this gap, we developed a novel homology-based algorithm, CGPFinder, which formalizes and automates the identification of clusters of gene paralogs (CGPs) by examining the physical distribution of individual gene members of families of paralogous genes across chromosomes. Application of CGPFinder to diverse mammalian genomes accurately identified CGPs for many well-known gene clusters in the human and mouse genomes (e.g., Hox, protocadherin, Siglec, and beta-globin gene clusters) as well as for 20 other mammalian genomes. Differences were due to the exclusion of non-homologous genes that have historically been considered parts of specific gene clusters, the inclusion or absence of one or more genes between the CGPs and their corresponding gene clusters, and the splitting of certain gene clusters into distinct CGPs. Finally, examination of human genes showing tissue-specific enhancement of their expression by CGPFinder identified members of several well-known gene clusters (e.g., cytochrome P450, aquaporins, and olfactory receptors) and revealed that they were unequally distributed across tissues. By formalizing and automating the identification of CGPs and of genes that are members of CGPs, CGPFinder will facilitate furthering our understanding of the evolutionary dynamics of genomic neighborhoods containing CGPs, their functional implications, and how they are associated with phenotypic diversity.

## Introduction

Gene duplications are among the most frequent types of mutational changes in genomes (Reams & Roth, 2015) and arguably the largest source of novel gene functions (Andersson & Hughes, 2009; Lynch & Conery, 2003; Zhang, 2003). Gene duplication can occur by many different mechanisms (Zhang, 2003), including transposition (Freeling et al., 2008), polyploidization (Carretero-Paulet & Fares, 2012; Grant, Cregan, & Shoemaker, 2000), and recombination (Krause & Pestka, 2015). Recombination-based gene duplication results in tandem gene duplication, in which the gene duplicates lie adjacent to each other and are physically linked on the chromosome (Glusman et al., 2000; Kawasaki & Weiss, 2003; Q. Wu & Maniatis, 1999). Chromosomal regions containing multiple homologs that have arisen through tandem gene duplication are common features of genomes, and are often described as paralogous gene clusters (Alam, Ain, Konno, Ho-Chen, & Soares, 2006; Krumlauf, 1992; MacLean et al., 2006; Martin, Freitas, Witt, & Christiansen, 2000; Noonan, Grimwood, Schmutz, Dickson, & Myers, 2004; Yagi, 2008).

Notable examples of paralogous gene clusters include the vertebrate protocadherin gene clusters (Noonan et al., 2004; Qiang Wu et al., 2001), the vertebrate and invertebrate olfactory receptor gene clusters (Hallem, Dahanukar, & Carlson, 2006; Niimura, Matsui, & Touhara, 2014; Niimura, 2009), the vertebrate natural killer cell receptor gene clusters (Kelley, Walter, & Trowsdale, 2005), and the Hox gene clusters found in one or more copies across metazoans (D. E. Ferrier & Holland, 2001; Glusman et al., 2000; Hoffman, Fernandez-Salguero, Gonzalez, & Mohrenweiser, 1995; Martin et al., 2000; Noonan et al., 2004). Many paralogous gene clusters contribute to traits that are highly variable, such as the composition of snake venom (Vonk et al., 2013), the architecture of animal body plan formation (Pendleton, Nagai, Murtha, & Ruddle, 1993), the olfactory repertoire (Glusman et al., 2000), or the immune response (Martin et al., 2000).

Given the many paralogous gene clusters from diverse gene families found in a wide diversity of organisms, the absence of a formal definition of what constitutes a “paralogous gene cluster” is surprising. The standard, informal definition that unites the known examples of paralogous gene clusters is that they represent groups of paralogous genes that are physically linked (Graham, 1995), although gene clusters sometimes also contain physically linked non-homologous genes (Krumlauf, 1994). As practical as this definition may be, it is subjective. For example, should two genes located right adjacent to each other on a chromosome be considered a cluster? Answering this question is challenging without considering the probability of observing two paralogs next to each other in the chromosome, which in turn requires knowledge of the number and distribution of paralogs in the genome as well as comparison of the intergenic distance between genes in the cluster with those in the rest of the chromosome.

An examination of the organization of the Hox gene cluster across diverse metazoans, which is often portrayed as a conserved, organized, and temporally and spatially clustered set of paralogous genes (Garcia-Fernàndez, 2005; Lemons & McGinnis, 2006; Pendleton et al., 1993), is a good case in point. Whereas genes in vertebrate Hox gene clusters are typically closely spaced and encoded on the same strand, Hox gene clusters in other animal phyla show striking differences in the number of genes that are members of the gene cluster, in their intergenic spacing, as well as in their general organization (Duboule, 2007). For example, the Hox gene cluster in sea urchin *Strongylocentrotus purpuratus* contains 2 non-Hox genes, and its constituent genes exhibit long intergenic distances and are encoded in both strands (Cameron et al., 2006; Duboule, 2007) (Fig. 1). In the fruit fly *Drosophila melanogaster*, the Hox “gene cluster” is actually composed by two distinct clusters; the ANT-C cluster, which contains 10 non-Hox genes and 7 Hox genes encoded in both strands (4 non-Hox genes and 6 Hox genes are in the Crick strand and the others in the Watson strand), and the BX-C cluster, which contains 3 Hox genes in the Crick strand and 1 non-Hox gene in the Watson strand (Fig. 1). In contrast, the HoxD gene cluster in the mouse *Mus musculus* – one of the four Hox clusters in this organism – is composed by nine contiguous homologous genes in the same strand (Fig. 1).

**Figure 1.**
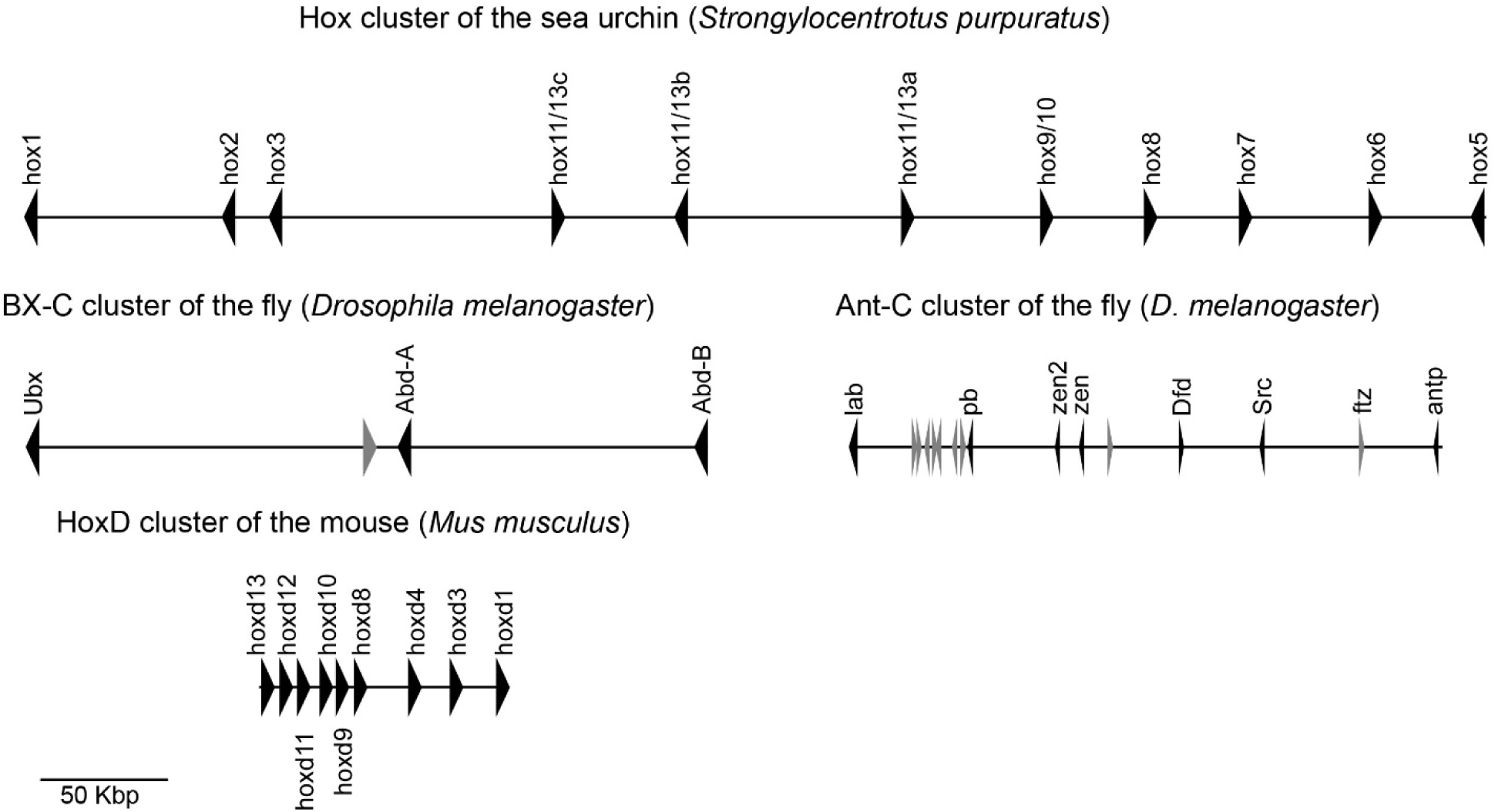
An illustration of the variation in the genomic organization of Hox “gene clusters” in three different animal species. The Hox gene cluster of the sea urchin (*Strongylocentrotus purpuratus*), the Bithorax complex (BX-C) and Antennapedia complex (Ant-C) clusters of the fruit fly (*Drosophila melanogaster*) and the HoxD cluster of the mouse (*Mus musculus*). Genes belonging to the Hox gene family are shown in black and intervening non-Hox genes in grey.

In this study, we propose a formal definition for a cluster of gene paralogs (CGP) that takes into account the sequence similarity of its members and their intergenic distances in the context of their chromosomal and genomic background to statistically assess whether neighboring paralogs form a CGP. We further implement our definition of CGP in CGPFinder, a computational tool for the identification of CGPs, and use it to examine the statistical validity of well-known gene clusters as well as explore the CGP landscape across different human tissues.

## Results

### Defining a Cluster of Gene Paralogs

We define a cluster of gene paralogs (CGP) in a given genome as a genomic region that contains a statistically significant higher number of paralogs from a specific gene family than the average background genomic region of the same length.

### The CGPFinder algorithm

To formally define and identify CGPs we developed the CGPFinder algorithm, which uses paralog sequence similarity and the density of the distribution of paralogs across the genome to statistically assess and demarcate the presence of CGPs in a genome. CGPFinder is written in Python and is freely available from https://github.com/biofilos/cgp_finder. Briefly, given a query reference protein sequence, or a set of homologous reference protein sequences, and a subject genome or set of genomes, CGPFinder uses the Blast algorithm (Altschul, Gish, Miller, Myers, & Lipman, 1990) to identify sequences that are statistically significantly similar (homologs) in the subject genome(s). The sets of paralogs identified on each of the chromosomes or genomic scaffolds of a given subject genome are considered candidate clusters.

Whether the genes in these candidate clusters comprise one or more genuine clusters of gene paralogs (CGPs) is evaluated and determined by the meanshift algorithm (Comaniciu, Meer, & Member, 2002). Specifically, the meanshift algorithm treats a parameter space as an empirical density function, and its objective is to find the region(s) of the parameter space with the highest density (or densities). In the context of the distribution of paralogs across a given chromosome, the meanshift algorithm identifies the genomic region(s) with the highest density (or densities) of paralogs of the protein reference sequence(s). Statistical assessment is performed by comparing the number of paralogs present in the genomic region(s) identified by the meanshift algorithm against an empirical distribution of paralogs.

Such an empirical distribution is obtained by counting the highest number of paralogs from any gene family contained in each of 1,000 randomly sampled genomic regions from the same genome of length equal to that of the cluster candidate from the meanshift step. Genomic regions with more paralogs than the genome-wide 95^th^ percentile of this empirical distribution are considered CGPs.

### CGPFinder recovers several well-known CGPs

To evaluate the performance of CGPFinder, we first examined whether it was able to identify a diverse set of previously characterized, well-known gene clusters in the human and mouse genomes. To capture as much sequence diversity as possible, we retrieved all the paralogs from each of 8 published gene clusters in either the mouse (*Mus musculus*) or in the human (*Homo sapiens*) genome and used them as a combined query in CGPFinder to identify CGPs in the same or in the other genome. The lists of mouse and human gene clusters used, and of the mouse and human genes used as queries for CGPFinder are described in Supplemental Table S1.

#### CGPFinder performance when searching the same genome

Using previously reported paralogous genes from 8 gene clusters found in the human and / or mouse genomes as queries (non-paralogous genes described as parts of these gene clusters were excluded from the blast query), CGPFinder correctly identified the CGPs in their corresponding genomes (Table 1). For example, all 5 genes reported to be part of the growth hormone gene cluster on the human chromosome 17 (Su Y. et al, 2000), all 26 protocadherin genes on the mouse chromosome 18 (Kohmura et al., 1998), all 11 HoxA genes in the human (chromosome 7; Krumlauf, 1994) and mouse (chromosome 6; Krumlauf, 1994) genomes, all 5 beta-globin genes in the human (chromosome 11; Levings & Bungert, 2002) and mouse (chromosome 5; Bulger et al., 1999; Weaver et al., 1981) genomes, and all 7 paralogous genes that are part of the luteinizing hormone beta (LHB) gene cluster on the human chromosome 19 (Hallast et al, 2008) were identified as statistically significant CGPs by CGPFinder.

**Table 1.** Comparison of eight human and mouse CGPs inferred by CGPFinder with the corresponding gene clusters previously described in the literature

| Query genes | Subject genome |  | Number of genes<br>in CGP<br>predicted by<br>CGPFinder | Notes/differences between predicted CGP and<br>previously described gene cluster | Reference |
| --- | --- | --- | --- | --- | --- |
|  | Species | Gene cluster<br>(chromosome) |  |  |  |
| The 11 HoxA genes<br>in chromosome 7<br>from *Homo sapiens* | *H. sapiens* | HoxA (7) | 11 | EVX1 was not used in the search because it is not<br>homologous to the genes in the query | (Krumlauf, 1992,<br>1994) |
|  | *H. sapiens* | HoxB (17) | 10 | In addition to the genes reported in the literature,<br>Hoxb13 was identified as part of the CGP |  |
|  | *H. sapiens* | HoxC (12) | 9 | none |  |
|  | *H. sapiens* | HoxD (2) | 9 | EVX2 was not used in the search because it is not<br>homologous to the genes in the query |  |
| The 11 HoxA genes<br>in chromosome 6<br>from *Mus musculus* | *M. musculus* | HoxA (6) | 11 | EVX1 was not used in the search because it is not<br>homologous to the genes in the query |  |
|  | *M. musculus* | HoxB (11) | 10 | In addition to the genes reported in the literature,<br>Hoxb13 was identified as part of the CGP |  |
|  | *M. musculus* | HoxC (15) | 9 | none |  |
|  | *M. musculus* | HoxD (2) | 9 | EVX2 was not used in the search because it is not homologous to the genes in the query |  |
| The 7 LHB genes in chromosome 19 from *Homo sapiens* | *H. sapiens* | LHB (19) | 7 | Genes RUVBL2 and NTF5 were not used in the search because they are not homologous to LHB genes | (Hallast, Saarela, Palotie, & Laan, 2008) |
| The 5 GH genes in chromosome 17 from *Homo sapiens* | *H. sapiens* | GH (17) | 5 | none | (Su Y. et al, 2000) |
| The 26 prolactin genes in chromosome 13 from *Mus musculus* | *M. musculus* | Prolactin (13) | 24 | In the 24-gene CGP, Prl2c1 was identified between Prl2a1 and Prl4a1, instead of the reported Plf. In addition to the reported paralogs, Gm3821 was also identified as part of the CGP. The remaining 3 genes (Prl2c4, Prl2c2 and Prl2c5) were identified as an independent cluster candidate but did not pass the statistical threshold to be considered a CGP | (Simmons et al., 2008) |
| The 7 galectin genes in chromosome 19 from *Homo sapiens* | *H. sapiens* | Galectins (19) | 4 / 3 | Genes previously reported as part of a single gene cluster were identified by CGPfinder as parts of two separate CGPs containing 4 and 3 galectin genes, respectively. LGALS17 was not included in the list of query genes and was not identified as part of the CGP because it is annotated as a pseudogene | (Than et al., 2009) |
| The 2 galectin genes<br>in chromosome 7<br>from *Mus musculus* | *M. musculus* | Galectins (7) | 2 | none | (Houzelstein et al.,<br>2004) |
| The 52<br>protocadherin genes<br>in chromosome 5<br>from *Homo sapiens* | *H. sapiens* | Protocadherins<br>(5) | 53 | In addition to the genes reported in the literature,<br>PCDHB16 was identified as part of the CGP | (Q. Wu &<br>Maniatis, 1999) |
| The 26<br>protocadherin genes<br>in chromosome 18<br>from *Mus musculus* | *M. musculus* | Protocadherins<br>(18) | 26 | none | (Kohmura et al.,<br>1998) |
| The 5 globin-beta<br>genes in<br>chromosome 11<br>from *Homo sapiens* | *H. sapiens* | Globins (11) | 5 | none | (Levings &<br>Bungert, 2002) |
| The 5 globin-beta<br>genes in<br>chromosome 7 from | *M. musculus* | Globins (7) | 5 | none | (Bulger et al.,<br>1999; Weaver et<br>al., 1981) |
| *Mus musculus* |  |  |  |  |  |
| The 9 SIGLEC genes in chromosome 19 from *Homo sapiens* | *H. sapiens* | SIGLECs (19) | 4 / 7 | Genes previously reported as part of a single gene cluster were identified as parts of two separate CGPs, containing 4 and 7 genes, respectively. In addition to the genes reported in the literature, the 4-gene CGP additionally contained SiglecL1, and the 7-gene CGP additionally contained loc105372490 | (Cao et al., 2009) |
| The 4 SIGLEC genes in chromosome 7 from *Mus musculus* | *M. musculus* | SIGLECs (7) | 7 | In addition to the genes reported in the literature, the CGP additionally contained 4931406B18Rik, Iglon5 and Vsig10L | (Angata, Margulies, Green, & Varki, 2004) |

CGPFinder also correctly identified known additional paralogous gene clusters in searches of the same genome (Table 1). For example, using all 11 HoxA genes from human chromosome 7 as a query, CGPFinder correctly identified CGPs corresponding the human HoxA, HoxB, HoxC, and HoxD clusters in the human genome; the same results were obtained for mouse (Table 1).

The correspondence between the identified CGPs and the previously described gene clusters was very good (Table 1) and the few observed differences fell into three categories. The first concerned differences associated with the exclusion of non-homologous genes that have historically been annotated as part of the gene cluster. For example, the EVX genes have historically been considered parts of mammalian Hox gene clusters (Krumlauf, 1994), but exhibit very low sequence similarity (14-28.5%) to the Hox genes and their inclusion simply reflects the knowledge that their function is associated with the function of the Hox gene cluster (Lemons & McGinnis, 2006). Because their sequence similarity to the Hox genes is not statistically significant, EVX genes were not included in the Hox CGPs identified by CGPFinder.

The second category included cases in which the CGPs contained one or a few additional genes not originally described as part of the gene cluster. For example, the HoxB CGPs in both the human (chromosome 17) and mouse (chromosome 11) genomes included Hoxb13, which was not originally described as part of the mammalian HoxB gene cluster (Krumlauf, 1994). Similarly, using the 52 protocadherin genes in human chromosome 5 as a query (Q. Wu & Maniatis, 1999), CGPFinder identified a CGP that additionally contained gene PCDHB16, whereas using the 4 Siglec genes on mouse chromosome 7 as a query (Kohmura et al., 1998), CGPFinder identified a 7-gene CGP that additionally contained genes 4931406B18Rik, Iglon5 and Vsig10L (Table 1).

The third and arguably most interesting category included cases in which CGPFinder split a previously described gene cluster into two distinct sub-clusters, which may or may not be both CGPs. For example, CGPFinder identified two distinct galectin CGPs, a 4-gene and a 3-gene one, instead of the single 7-gene cluster previously reported to reside on human chromosome 19 (Than et al., 2009). CGPFinder identified two separate CGPs because the average intergenic distances of the genes in the 3-gene and 4-gene CGPs are 13 Kbp and 38 Kbp, respectively, which are significantly smaller than the 790 Kbp that separates the two CGPs. Similarly, genes previously reported as part of a single 9-gene Siglec cluster on human chromosome 19 (Cao et al., 2009) were identified by CGPFinder as parts of two separate CGPs, containing 4 and 7 genes (with intergenic distances of 38 Kbp and 29 Kbp), respectively, separated by 142 Kbp. In addition to the genes reported in the literature, the 4-gene CGP additionally contained the SiglecL1 gene, and the 7-gene CGP additionally contained the loc105372490 gene. Interestingly, the two CGPs directly correspond with the A and B sub-clusters that resulted from an inverse duplication of the CD33rSiglec cluster in eutherian mammals (Cao et al., 2009). Finally, using the 26 genes in the prolactin cluster on mouse chromosome 13 (Simmons, Rawn, Davies, Hughes, & Cross, 2008) as a query, CGPFinder identified a single 24-gene CGP (Table 1). An additional candidate gene cluster located 13.7 Mbp away and comprised of the genes Prl2c3, Prl2c2 and Prl2c5 was also identified, but it was below the 95^th^ percentile of the empirical distribution of paralogs for a genomic region of that size and was not recognized as a CGP.

#### CGPFinder performance when searching different genomes

Given that CGPFinder correctly identified several well-known gene clusters in searches of the same genome, including known additional paralogous gene clusters (Table 1), we next sought to examine the performance of CGPFinder on the human genome when using gene queries from the mouse genome (Table 2) as well as on the mouse genome when using gene queries from the human genome (Table 3).

**Table 2.** Comparison of mouse CGPs using human queries inferred by CGPFinder with gene clusters previously described in the literature

| Query genes | Subject genome |  | Number of genes in CGP predicted by CGPFinder | Notes/differences between predicted CGP and previously described gene cluster | Reference |
| --- | --- | --- | --- | --- | --- |
|  | Species | Gene cluster (chromosome) |  |  |  |
| The 11 HoxA genes in chromosome 7 from *H. sapiens* | *M. musculus* | HoxA (6) | 11 | EVX1 was not used in the search because it is not homologous to the genes in the query | (Krumlauf, 1992, 1994) |
|  | *M. musculus* | HoxB (11) | 10 | In addition to the genes reported in the literature, Hoxb13 was identified as part of the CGP |  |
|  | *M. musculus* | HoxC (15) | 9 | none |  |
|  | *M. musculus* | HoxD (2) | 9 | EVX2 was not used in the search because it is not homologous to the genes in the query |  |
| The 7 galectin genes in chromosome 19 from *H. sapiens* | *M. musculus* | Galectins (7) | 2 | none | (Houzelstein et al., 2004) |
| The 52 protocadherin | *M. musculus* | Protocadherins (18) | 24 | Pcdh1 and Pcdh12 were not found (these two genes were found as part of the CGP when mouse genes | (Kohmura et al., 1998) |
| genes in<br>chromosome 5<br>from *H. sapiens* |  |  |  | were used as the query) |  |
| The 5 globin-beta<br>genes in<br>chromosome 11<br>from *H. sapiens* | *M. musculus* | Globins (7) | 5 | none | (Bulger et al.,<br>1999; Weaver et<br>al., 1981) |
| The 9 SIGLEC<br>genes in<br>chromosome 19<br>from *H. sapiens* | *M. musculus* | SIGLECs (7) | 6 | In addition to the genes reported in the literature,<br>4931406B18Rik and Vsig10L were identified as<br>part of the CGP | (Angata et al.,<br>2004) |

**Table 3.** Comparison of human CGPs using mouse queries inferred by CGPFinder with gene clusters previously described in the literature

| Query genes | Subject genome |  | Number of genes in the cluster predicted by CGPFinder | Notes/differences between predicted CGP and previously described gene cluster | Reference |
| --- | --- | --- | --- | --- | --- |
|  | Species | Cluster (chromosome) |  |  |  |
| The 11 HoxA genes in chromosome 6 from *M. musculus* | *H. sapiens* | HoxA (7) | 11 | EVX1 was not used in the search because it is not homologous to the genes in the query | (Krumlauf, 1992, 1994) |
|  | *H. sapiens* | HoxB (17) | 10 | In addition to the genes reported in the literature, Hoxb13 was identified as part of the CGP |  |
|  | *H. sapiens* | HoxC (12) | 9 | none |  |
|  | *H. sapiens* | HoxD (2) | 9 | EVX2 was not used in the search because it is not homologous to the genes in the query |  |
| The 2 galectin genes in chromosome 7 from *M. musculus* | *H. sapiens* | Galectins (19) | 4 / 3 | Genes previously reported as part of a single gene cluster were identified by CGPFinder as parts of two separate CGPs containing 4 and 3 galectin genes, respectively | (Than et al., 2009) |
| The 26 protocadherin genes in chromosome 18 from *M. musculus* | *H. sapiens* | Protocadherins (5) | 55 | In addition to the genes reported in the literature, PCDHB16, PCDH1, PCDH12 were identified as part of the CGP | (Q. Wu & Maniatis, 1999) |
| The 5 globin-beta genes in chromosome 7 from *M. musculus* | *H. sapiens* | Globins (11) | 5 | none | (Levings & Bungert, 2002) |
| The 4 SIGLEC genes in chromosome 7 from *M. musculus* | *H. sapiens* | SIGLECs (19) | 5 / 7 | Genes previously reported as part of a single gene cluster were identified as parts of two separate CGPs, containing 5 and 7 genes, respectively. In addition to the genes reported in the literature, the 5-gene CGP additionally contained SiglecL1 and VSIG10L, and the 7-gene CGP additionally contained<br>loc105372490 | (Cao et al., 2009) |

In general, CGPFinder correctly identified CGPs for several well-known gene clusters in the human and mouse genomes. For example, using the 11 HoxA genes from the human genome as a query, CGPFinder correctly identified a CGP corresponding to the mouse HoxA gene cluster (Table 2); similarly, using the 11 mouse HoxA genes as a query, CGPFinder identified the human HoxA gene cluster (Table 3). The same was true for the mouse galectin and beta-globin gene clusters, which were identified using their human homologs (Table 2), as well as for the human beta-globin gene cluster, which was identified by CGPFinder using mouse homologs (Table 3). CGPFinder also correctly identified known additional paralogous gene clusters in a given genome. For example, using all 11 HoxA genes from human chromosome 7, CGPFinder correctly identified CGPs corresponding to the HoxA, HoxB, HoxC, and HoxD clusters in the mouse genome (Table 2). Likewise, using the mouse HoxA genes as a query, CGPFinder correctly identified CGPs corresponding to the human HoxA, HoxB, HoxC and HoxD gene clusters (Table 3).

Similarly to the results of the performance of CGPFinder when searching the same genome, the differences between the inferred CGPs and the previously described gene clusters when searching other genomes fell into three categories. The first category concerned differences associated with the exclusion of non-homologous genes that have historically been considered to be parts of specific gene clusters, the second included single or a few gene differences between the CGPs and their corresponding gene clusters, and the third cases in which a previously described gene cluster was split by CGPFinder into two distinct CGPs (Tables 2 and 3).

The most conspicuous differences between the identified CGPs and the previously described gene clusters were observed in the mouse and human Siglec gene clusters. Specifically, using the 9 Siglec genes on human chromosome 19 as a query, CGPFinder identified a 6-gene CGP on mouse chromosome 7 that contained two additional genes (4931406B18Rik and Vsig10L) in addition to those previously described for the mouse Siglec gene cluster (Angata et al., 2004). Furthermore, using the 4 Siglec genes on mouse chromosome 7 as a query, CGPFinder identified two separate CGPs 69.0 Kbp away from each other, containing 5 and 7 genes (with average intergenic distances of 44.5 Kbp and 29.4 Kbp), respectively, that correspond to the A and B sub-clusters previously identified by Cao and co-workers (2009; Table 2). In addition to the genes previously reported (Cao et al., 2009), the 5-gene CGP additionally contained SiglecL1 and VSIG10L, and the 7-gene CGP additionally contained loc105372490 (Table 3). Interestingly, the human gene VSIG10L was identified as part of this CGP only when using mouse Siglec genes as queries (Tables 1 and 3). This is because VSIG10L shows statistically significant sequence similarity only to the mouse gene Siglecg but not to any human Siglec genes.

### Identifying CGPs across placental mammals

Given that CGPFinder performed very well in recovering several well-known CGPs in the human and mouse genomes, we next used all the paralogs from each of 6 published gene clusters in the human genome and used them as a combined query in CGPFinder (Supplemental Table S1) to identify CGPs from the same six gene families (galectin, Hox, beta-globin, Siglec and protocadherin) in 20 other high quality mammalian genomes (Fig. 2, Supplemental Table S2).

**Figure 2.**
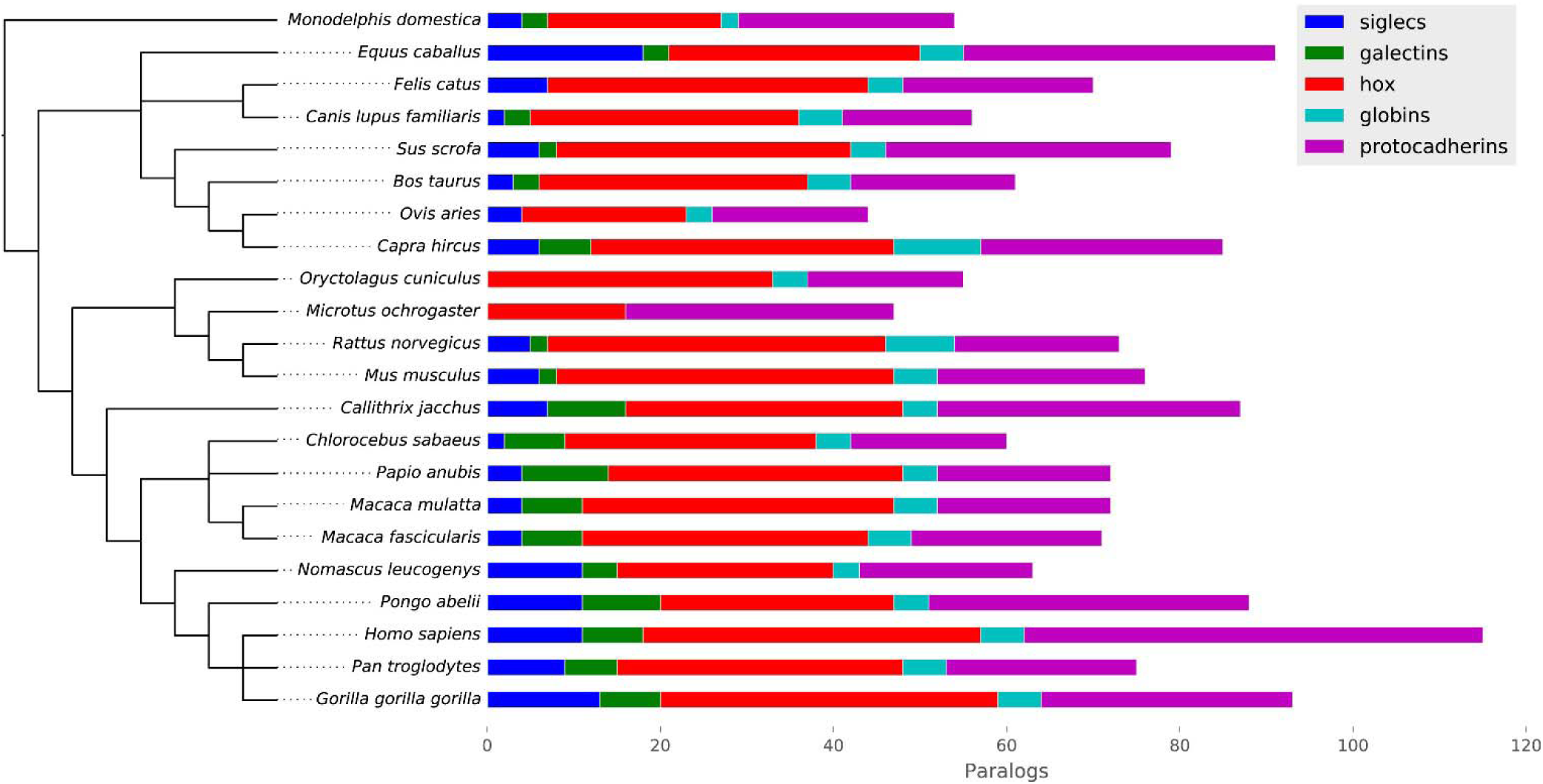
Distribution of CGPs in five selected gene families across 22 mammalian genomes. For gene families with more than one CGP in a given organism, numbers of paralogs reflect the total number found in all CGPs associated with that gene family.

Overall, CGPFinder identified all the CGPs that are expected to be present and conserved in these 20 mammalian genomes (Supplemental Table S2). For example, running CGPFinder with the HoxA genes from human as a query identified four Hox clusters in all the studied genomes, with the exceptions of HoxA and HoxB in vole (*Microtus ochrogaster*), HoxB in orangutan (*Pongo abelii*), and HoxC in opossum (*Monodelphis domestica*) (Supplemental Table S2). The reason for these exceptions is that these clusters are present in non-assembled scaffolds that are not part of the standard genome assemblies provided by GenBank. Similarly, a CGP for the protocadherin cluster was identified in all species using the genes from the protocadherin cluster in human chromosome 5 as a query. However, more than one CPGs in the same chromosome were identified in cow (*Bos taurus*), gorilla (*Gorilla gorilla*), dog (*Canis lupus*), sheep (*Ovis aries*) and pig, and protocadherin CGPs in more than one chromosome were identified in gorilla, human, and pig (Supplemental Table S2).

Using the human beta-globins as the query, CGPFinder also identified both alpha-and beta-globin CGPs in all species, except in vole where only the alpha-globin CGP was found, and in horse (*Equus caballus*) and rabbit (*Oryctolagus cuniculus*) where only the beta-globin CGP was found (alpha-and beta-globin CGPs were differentiated by constructing their phylogeny; see methods). The number of genes contained in the beta-globin CGP varied across species (Fig. 2, Supplemental Tables S2 and S3). The largest beta-globin CGP was found in goat (*Capra hircus*) and stemmed from a previously reported beta-globin cluster duplication (Hardies, Edgell, & Hutchison, 1984), followed by the rat (*Rattus norvegicus*) 8-gene CGP. Most species contained either 5-, 4-, or 3-gene CGPs, but opossum (*Monodelphis domestica*) contained a 2-gene CGP. This variation is likely due to both paralog gain and loss as well as errors in annotation.

Running CGPFinder with the 52 reported genes from the human protocadherin cluster (Q. Wu & Maniatis, 1999) as a query identified one protocadherin CGP per species, with the number of paralogs per CGP ranging from 15 in dog (*Canis lupus familiaris*), to 53 in human (Supplemental Table S2). The only exception was pig, where a 13-gene CGP (average intergenic distance of 13.1 Kbp) and a 20-gene CGP (average intergenic distance of 16.0 Kbp) were found on the same chromosome but separated by 213 Kbp.

The galectin CGP in all species was identified by CGPFinder using the 7-gene human galectin cluster reported by Than and collaborators (2009) as a query. Orthologous galectin CGPs were extracted from all the galectin CGPs using a phylogenetic tree rooted on the clade containing the galectins LGALS1 and LGALS2, following a previously reported galectin phylogeny (Houzelstein et al., 2004) (Fig. 2, Supplemental Tables S2 and S3). No orthologous CGPs were found in cat (*Felis catus*), vole, rabbit and sheep, although vole and cat contained other homologous galectin CGPs. One orthologous 3-gene CGP was found in cow, dog and horse, whereas mouse, rat and pig contained one orthologous 2-gene CGP. All other mammals contained two orthologous galectin CGPs containing 2-3, and 3-7 genes respectively (Supplemental Table S2).

Using the protein sequences from the previously reported Siglec cluster in human (Cao et al., 2009), CGPFinder identified a Siglec CGP in cow, dog, goat, green monkey (*Chlorocebus sabaeus*), horse, cat, rhesus macaque (*Macaca mulatta*), mouse, sheep, chimpanzee (*Pan troglodytes*), baboon (*Papio Anubis*), orangutan, rat, and pig (Supplemental Table S2). The number of paralogs in these CGPs ranged from 2 in dog and green monkey, to 11 in orangutan and 18 in horse. No Siglec CGP was found in rabbit and vole. Two Siglec CGPs were found in marmoset (*Callithrix jacchus*), gorilla (*Gorilla gorilla*), crab-eating macaque (*Macaca fascicularis*), human, and gibbon (*Nomascus leucogenys*). In all these cases, the number of genes in the first (ordered by genomic coordinates) CGP (ranging between 2 to 6 paralogs) was smaller or equal to the number of genes in the second CGP (2 to 9 paralogs) (Fig. 2, Supplemental Table S2 and S3).

Finally, CGPFinder accurately identified CGPs for clusters that are known to be taxonomically-restricted to certain lineages. For example, using the genes from the human growth hormone and LHB clusters, CGPFinder identified CGPs for growth hormone and LHB in primate genomes (Supplemental Table S2), consistent with the previously described primate-specific presence of these gene clusters (Hallast et al., 2008; Su Y. et al, 2000). However, a small CGP of two LHB paralogs was identified in cow and horse; interestingly, the duplications that led to the formation of these CGPs appear to have been independent of the duplications that led to the formation of the primate LHB cluster. Similarly, using the genes from the prolactin cluster in mouse as a query, CGPFinder was able to identify CGPs for previously described prolactin clusters in certain rodent (rat and vole) and bovid species (cow) (Simmons et al., 2008; Wallis, 1991) (Supplemental Table S2). Among the bovids, a previously unreported 12-gene prolactin CGP was identified in goat (*Capra hircus*).

## What fraction of genes showing enhanced expression in human tissues is clustered?

To illustrate the potential of CGPFinder as a tool for furthering our understanding of the function of CGPs in the human genome, we used all 4,866 genes showing tissue-specific enhancement of their expression in 32 human tissues and organs (enhanced genes) (Uhlen et al., 2015) as individual queries in CGPFinder to examine whether they were clustered or not. On average, 25% of enhanced genes was part of CGPs. However, clustered genes were unequally distributed across tissues, with the percentage of clustered genes per tissue ranging from 56% (49/88) in the appendix, to 12% (74/597) in the cerebral cortex (Fig. 3).

**Figure 3.**
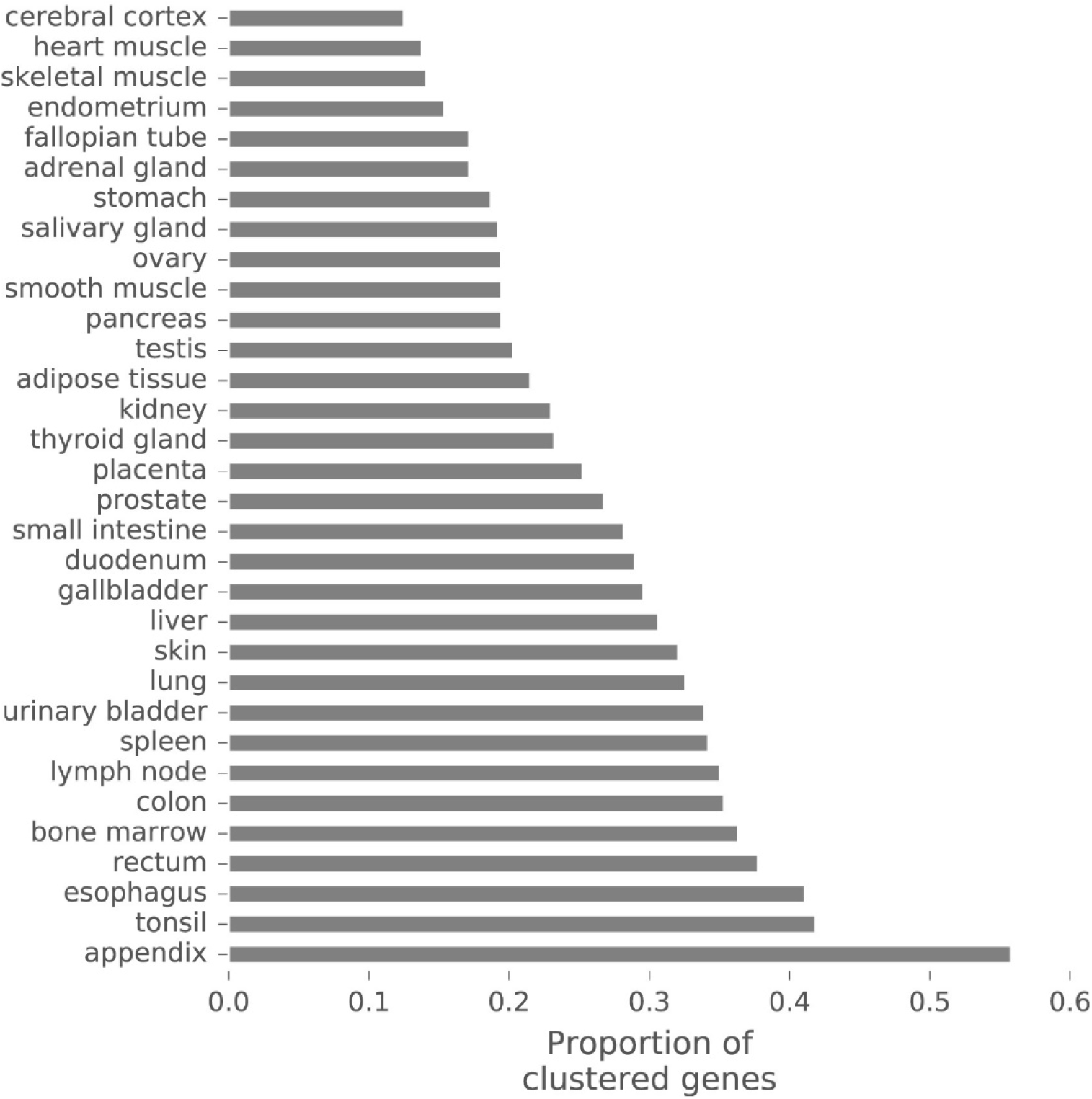
Relative abundance of clustered genes (i.e., genes contained in CGPs) showing tissue-specific enhancement of their expression across 32 human tissues and organs.

Examination of the enhanced genes that reside within CGPs identified members of several well-known gene clusters. In general, gene families that are known to be parts of gene clusters like olfactory receptors (Niimura, Niimura, & Y, 2009), phospholipase A (Tischfield et al., 1996), golgins (Locke et al., 2003), histones (Albig, Kioschis, Poustka, Meergans, & Doenecke, 1997), cytochrome P450s (Hoffman et al., 1995), aquaporins (Finn, Chauvigné, Hlidberg, Cutler, & Cerdà, 2014), myosin chains (Weiss et al., 1999), Hox genes (Krumlauf, 1992), and protocadherins (Q. Wu & Maniatis, 1999), were enhanced in different tissues (Supplemental Table S4). For example, 9 of the 49 clustered genes in the appendix are chemokine ligands or receptors, 6 are leukocyte immunoglobulin-like receptors, and 4 are SIGLECs. Similarly, 20 of the 75 clustered genes in the bone marrow are annotated as part of the histone cluster 1, 32 of 72 clustered genes in the lymph node are involved in the immune system (e.g., chemokines, MHC, T-cell and B-cell associated), and 6 of 12 clustered genes in the endometrium belong to the Hox gene family (clusters A and D). Finally, in the placenta, clustered genes encode for proteins important in tissue structure and remodeling like metallopeptidases (2 of 41 clustered genes), collagen (3 genes) and gap junction proteins (3 genes), as well as immunity related genes (one interleukin receptor and one immunoglobulin receptor) (Supplemental Table S4).

## Discussion

From the classic globin and Hox gene clusters (Efstratiadis, 1980; Krumlauf, 1994) to the more recently described venom-related gene clusters in snakes (Ikeda et al., 2010; Vonk et al., 2013), the study of paralogous gene families that are physically linked on chromosomes has greatly enhanced our understanding of physiology, development, and the genetic basis of phenotypic diversity (Hallem et al., 2006; Holland & Garcia-Fernàndez, 1996; Rawn & Cross, 2008; Whittington et al., 2008). In the post-genomic era, however, the fact that such gene clusters are still defined arbitrarily and in different ways in each gene family or genome makes attempts for the kinds of comparative analyses required to understand the dynamics of gene cluster evolution and function problematic.

Our formal definition and identification of clusters of gene paralogs (CGPs; implemented as CGPFinder) through a statistical approach that takes into account both intergenic distance and homology solves this problem. This approach not only enables the comparison of the genomic arrangement of well-known clusters in different model organisms, but also the discovery of novel gene clusters or novel arrangements in genomes. For example, by using previously known sequences of the prolactin gene cluster in mouse, CGPFinder was able to find novel prolactin CGPs in goat and sheep (Supplemental data; Supplemental Table S2), providing further evidence supporting the independent evolution of prolactin clusters in ruminants and rodents (Miller & Eberhardt, 1983). Thus, coupled with robust phylogenetic analysis at the gene and species levels, CGPFinder could be very useful for distinguishing between different types of clusters (e.g., primary, secondary, and independently evolved) comprised of paralogous genes and the evolutionary history of their assemblies (Ferrier, 2016).

Using a formal definition for CGPs also has the potential to help identify interesting gene arrangement and orientation features in clusters and inform our understanding of the evolutionary steps that explain their current assembly. For example, according to CGPFinder, the single Siglec gene cluster on the human chromosome 19 is actually comprised of 2 distinct CGPs (Tables 1 and 3). Interestingly, the two inferred CGPs correspond precisely to the two sub-clusters (A and B) that Cao and co-workers previously identified and inferred to have been generated through a large-scale inverse duplication of the ancestral Siglec locus in a vertebrate ancestor (Cao et al., 2009). Such inversions can help stabilize the size of a gene cluster by reducing the effectiveness of recombination to add or remove additional gene duplicates to the existing cluster (Cao et al., 2009; Passananti, Davies, Ford, & Fried, 1987).

Availability of a formal definition and means of characterizing CGPs will also facilitate efforts to understand the functional implications of clustering. For example, at a broad level, CGPFinder can be used to understand whether tissue-specific expression of clustered genes is evenly distributed across human tissues and organs (Fig. 3), which in turn could be associated with the types and functions of genes expected to function in them (e.g., many secreted proteins are known to be expressed in tissues such as the liver and salivary glands) (Uhlen et al., 2015). Additionally, given that CGPFinder can simultaneously analyze the genomes of multiple species, a formal definition enables the investigation of the types of functional categories of genes (e.g., immunity, metabolism) that tend to be clustered in the genomes of organisms from diverse lineages and which potentially could be implicated in the generation of lineage-specific interesting phenotypes.

Finally, a formal definition for CGPs has the potential to greatly aid in understanding the relationship between the specific genomic organization of a given gene cluster with its mechanism of regulation. For example, using a diverse and rich body of data on animal Hox gene clusters, Duboule (2007) has argued that Hox gene cluster organization in different animals (Fig. 1) has strong implications for how the activity of Hox genes is regulated in these organisms. The first step toward answering this question, not just for the animal Hox gene clusters, but for the wide diversity of gene families forming gene clusters, is the availability of a clear, precise, unambiguous and easy to implement definition, such as the one provided by this study.

## Methods

### Gene annotation

Genome annotation for all the mammalian species with assembled chromosomes available from the NCBI was downloaded from ftp.ncbi.nlm.nih.gov/genomes/ (Supplemental Table S5). Genes were annotated using CDS (coding sequence) features from their annotation GenBank files. If two or more CDS features had the same start or end coordinate, the longest gene was selected. If two genes in the same strand had overlapping coordinates but different start coordinates, the gene with the start coordinates downstream from the other gene was removed.

### CGPFinder validation on previously described (reference) CGPs

Representative protein sequences from the prolactin, growth hormone, Hox, galectin, luteinizing hormone beta (LHB), beta-globin, Siglecs, galectin, and protocadherin gene clusters were downloaded from the NCBI. Gene family cluster analysis was performed for each downloaded set of sequences using the CGPFinder algorithm described below.

### Phylogenetic analysis of orthologous CGPs

In order to identify orthologous CGPs, phylogenetic trees were built using the protein sequence from all the CGPs in a gene family. The best substitution model was selected using ProtTest (Darriba, Taboada, Doallo, & Posada, 2011). The phylogenetic tree was estimated using the rapid bootstrap function implemented in RAxML, version 8.2.4 (Stamatakis, 2006), and 100 bootstrap pseudoreplicates were performed. The tree was rooted using evolutionary information about each family, and the leaves containing genes from known CGPs in human and mouse were used to identify the clade with CGP orthologs (selected clade). Given that CGPs in human and mouse could have experienced gene deletions or accelerated rates of evolution, sister branches of the selected clade were also inspected for the presence of members of the known CGPs. The set of CGP orthologs was defined as all the CGPs with all their genes included in the selected clade.

### CGPFinder programing environment

CGPFinder was coded in Python 3.4.3 using the Pandas library v0.16.2 for table manipulation, Numpy v1.9.2 (Oliphant, 2007) for numerical arrays manipulation, BioPython v1.65 (Cock et al., 2009) for sequence manipulation and communication with the NCBI servers, scikit-learn v0.16.1 (Pedregosa et al., 2011) for statistical analysis, and matplotlib v1.4.3 (Hunter, 2007) and Bokeh v.0.9.2 for graphics generation. CGPFinder is freely available from https://github.com/biofilos/cgp_finder.

### Assessing the degree of clustered of genes with human tissue-enhanced expression

The “RNA gene data” expression dataset was downloaded from the Protein Atlas (tissue dataset) (Uhlen et al., 2015), and genes that were tissue-enhanced were selected. Protein sequences from these selected genes were downloaded using the BioMart API from Ensembl. In cases where several protein sequences were mapped to the same gene identifier, only the longest protein sequence was used. CGPFinder was run for each of the selected genes. To evaluate whether a given gene was a member of a CGP, its coordinates were contrasted with those of all the clustered genes found by CGPFinder. If the coordinates of a selected gene overlapped with those of a clustered gene, the gene was annotated as clustered.

### Gene family cluster analysis (CGPFinder algorithm)

Each query protein or protein set was used as a query for blastp (E-value 0.0001) against all the proteomes of the species under study. Blast hits that were either more than three times the length of the query or less than one third the length of the query were removed (length ratio less than 0.3). The set of blastp hits in each chromosome (pre-clusters) was processed using the meanshift algorithm as implemented in scikit-learn (Pedregosa et al., 2011). Since the meanshift algorithm is implemented to identify clusters in more than one dimension, the start coordinates of the pre-clusters were considered to be the x coordinates and the y coordinates were set to zero. The mean plus the standard deviation of all intergenic distances in the chromosome where a given pre-cluster resides were used as the bandwidth parameter for the meanshift implementation. In the meanshift step, the pre-clusters can be ignored, subdivided, or confirmed as proto-clusters. Its output consists of a set of proto-clusters per chromosome.

In order to extract clusters that have more paralogs than expected in the genome, 1,000 random regions of the length of each proto-cluster were taken from the genome for the following analysis: an all vs. all blastp search of the complete set of proteomes under study was used to extract the paralogs of the genes in the region, and the maximum number of hits was considered the maximum number of paralogs. Proto-clusters containing more paralogs than the 95^th^ percentile from the genome-wide sample set were considered clusters of gene paralogs (CGP).

## ACKNOWLEDGMENTS

We thank members of the Rokas lab for critical feedback on the work described in this study. This work was conducted in part using the resources of the Advanced Computing Center for Research and Education at Vanderbilt University. JFO was supported by the Graduate Program in Biological Sciences at Vanderbilt University. Research on this project was supported in part by the March of Dimes through the March of Dimes Prematurity Research Center Ohio Collaborative and by the National Science Foundation (DEB-1442113 to A.R.).

